# Interspecific differences in second-site modifiers reshape dominance in a Mendelian trait by altering environmental plasticity

**DOI:** 10.64898/2026.10.06.757123

**Authors:** Yuichi Fukutomi, Alexandra Phillips-Garcia, Jianwei Zhang, Artyom Kopp

## Abstract

Dominance is an important and evolvable property of the genotype–phenotype map. A longstanding debate in evolutionary genetics concerns whether dominance is determined primarily by intrinsic properties of alleles at the focal locus or by modifiers elsewhere in the genome. Theoretical models show that second-site modifiers of dominance can evolve under some conditions. However, direct empirical tests of these models are rare because they require holding the focal heterozygous genotype constant while changing the rest of the genome. In the *Drosophila montium* species subgroup, female pigmentation is a Mendelian trait controlled by a single gene, *POU domain motif 3* (*pdm3*), but the dominance relationships between *pdm3* alleles differ among species. Using reciprocal introgressions among three closely related species, we compared females carrying the same Light/Dark *pdm3* allele combinations in different homozygous or heterozygous genomic backgrounds and across different temperatures. In both species-pair comparisons, changing genomic background significantly altered the phenotype of identical *pdm3* heterozygotes, demonstrating the existence of second-site modifiers of dominance. The effects of dominance modifiers depend on the particular pair of *pdm3* alleles and on rearing temperature. These results show that dominance in this system is a genotype-by-genotype-by-environment trait, rather than a fixed attribute of individual alleles. The finding that second-site modifiers can alter dominance by reshaping reaction norms provides a developmental mechanism that links the evolution of dominance to phenotypic plasticity.

**Significance:** The causes of allelic dominance, and the mechanisms of its evolution, have been debated since the early work of Fisher, Wright, and Haldane 100 years ago. Recent theoretical models predict that second-site modifiers of dominance can evolve under realistic conditions, but empirical evidence remains scarce and contradictory. This study demonstrates the existence of modifiers that change dominance relationships at a Mendelian locus that controls interspecific differences in *Drosophila* pigmentation by testing the same pairs of focal alleles in the genomic backgrounds of different species. These modifiers act by altering phenotypic plasticity, providing a previously unappreciated developmental mechanism for the evolution of dominance.

## Introduction

Dominance - the extent to which the phenotypic effect of an allele is expressed in heterozygotes - is a key part of the relationship between genotype and phenotype. By affecting the efficiency of selection, dominance influences the evolutionary fate of mutations, shaping their probabilities of persistence and the amount of genetic variation maintained in populations (Billiard et al. 2021; Di and Lohmueller 2024). The causes of dominance have been debated since the earliest days of population genetics. Fisher (1928) proposed that dominance evolves through natural selection on second-site modifiers, so that the phenotypic relationship between alleles at a focal locus depends on alleles elsewhere in the genome. Wright (1929) argued instead that dominance mainly reflects the physiological properties of gene products, without requiring second-site modifiers or selection for dominance. Haldane (1930) likewise treated dominance as a property of alleles at the focal locus, but proposed that natural selection could favor alleles with greater dominance. Wright and Haldane argued that selection for second-site modifiers would typically be too weak to drive their evolution because their epistatic effects are contingent on rare heterozygous genotypes (Wright 1929; Haldane 1930). However, recent models show that such modifiers can evolve when heterozygotes at the focal locus are sufficiently common – for example, under balancing, frequency-dependent, or sexually antagonistic selection (Bourguet 1999; Peischl and Bürger 2008; Billiard and Castric 2011; Llaurens et al 2015; Spencer and Priest 2016; Li and Bank 2024).

Empirical studies show that dominance is evolvable and context-dependent (Billiard et al. 2021; Di and Lohmueller 2024; Grieshop et al. 2024; Li and Bank 2024). For the same heterozygous genotype, dominance can differ between sexes (Barson et al. 2015; Grieshop and Arnqvist 2018; Pearse et al. 2019) or environments (Posavi et al. 2014; Karageorgi et al. 2025). Dominance can also vary among populations or species (Stewart and Lees 1987; Bourguet et al. 1996; Nijhout 2003; Rosenblum et al 2010; Le Poul et al 2014; Fukutomi et al 2026), making it possible to test whether dominance evolves because of changes at the focal alleles themselves or at second-site modifiers. Empirical evidence remains both sparse and contradictory. Some cases point to focal-allele effects: opposite dominance of body color in two lizard species is caused by distinct coding mutations in the *Mc1r* gene (Rosenblum et al. 2010), and reversed dominance for heterodichogamy in two genera of Juglandaceae is associated with independently evolved haplotypes at the *GFAFL* gene (Groh et al. 2025). At plant self-incompatibility loci, dominance is controlled by small RNAs located within the focal S-haplotype (Tarutani et al. 2010; Durand et al. 2014), while in moths, melanic alleles retain their dominance after introgression into different genetic backgrounds (West, 1977; Mikkola, 1984). Conversely, several studies show that genetic background can alter dominance, as in the case of butterfly wing color loci (Naisbit et al. 2003; Nijhout 2003), *Tetrahymena* mating type alleles (Phadke et al. 2014), or a transposable element insertion affecting oxidative stress response in *Drosophila* (Guio and González 2015). To date, reciprocal experimental tests that hold the focal allele pair constant while changing the genomic background in a natural system where dominance has evolved remain very limited.

A major unresolved question is *how* second-site modifiers can alter dominance at the focal locus. One possibility is that dominance evolves as a consequence of changes in phenotypic plasticity. Classical studies showed that selection for developmental robustness can simultaneously modify dominance, suggesting environmental canalization as an indirect route to dominance evolution (Rendel 1959; Sheppard and Ford 1966; Belt and Burnet 1972). More generally, nonlinear genotype–phenotype maps can make heterozygotes either more robust or more plastic than homozygotes, depending on their position on the underlying developmental response curve (Green et al. 2017; Klingenberg 2019; Billiard et al. 2021; Grieshop et al. 2024; Fukutomi et al. 2026; David et al. 2026). Thus, the evolution of plasticity and dominance may represent two consequences of the same underlying developmental change. This mechanism could also make the evolution of second-site dominance modifiers more likely, because modifiers that affect environmental sensitivity can be selected through their effects in both homozygotes and heterozygotes at the focal locus.

The *Drosophila montium* species subgroup provides an opportunity to test whether the evolution of dominance reflects changes at the focal locus, second-site modifiers, or their interaction with the environment. Many species in this clade show a single-locus, female-limited abdominal color dimorphism (Yassin et al. 2016) (Figure 1). Dark color is fully dominant in most species, but two distantly related species with dominant light pigmentation have been described (Ohnishi and Watanabe 1985; Yassin et al. 2016; Prigent et al. 2020). In all studied species, this Mendelian dimorphism is controlled by the *POU domain motif 3* (*pdm3*) gene, which encodes a transcription factor that represses dark pigmentation (Yassin et al. 2016; Fukutomi et al. 2026). The dominance reversal is also coupled to thermal plasticity: in dark-dominant species, the Dark *pdm3* allele is fully dominant at all temperatures, whereas in light-dominant species the Light *pdm3* allele becomes increasingly dominant as temperature increases (David et al. 2026; Fukutomi et al. 2026).

**Figure 1.**
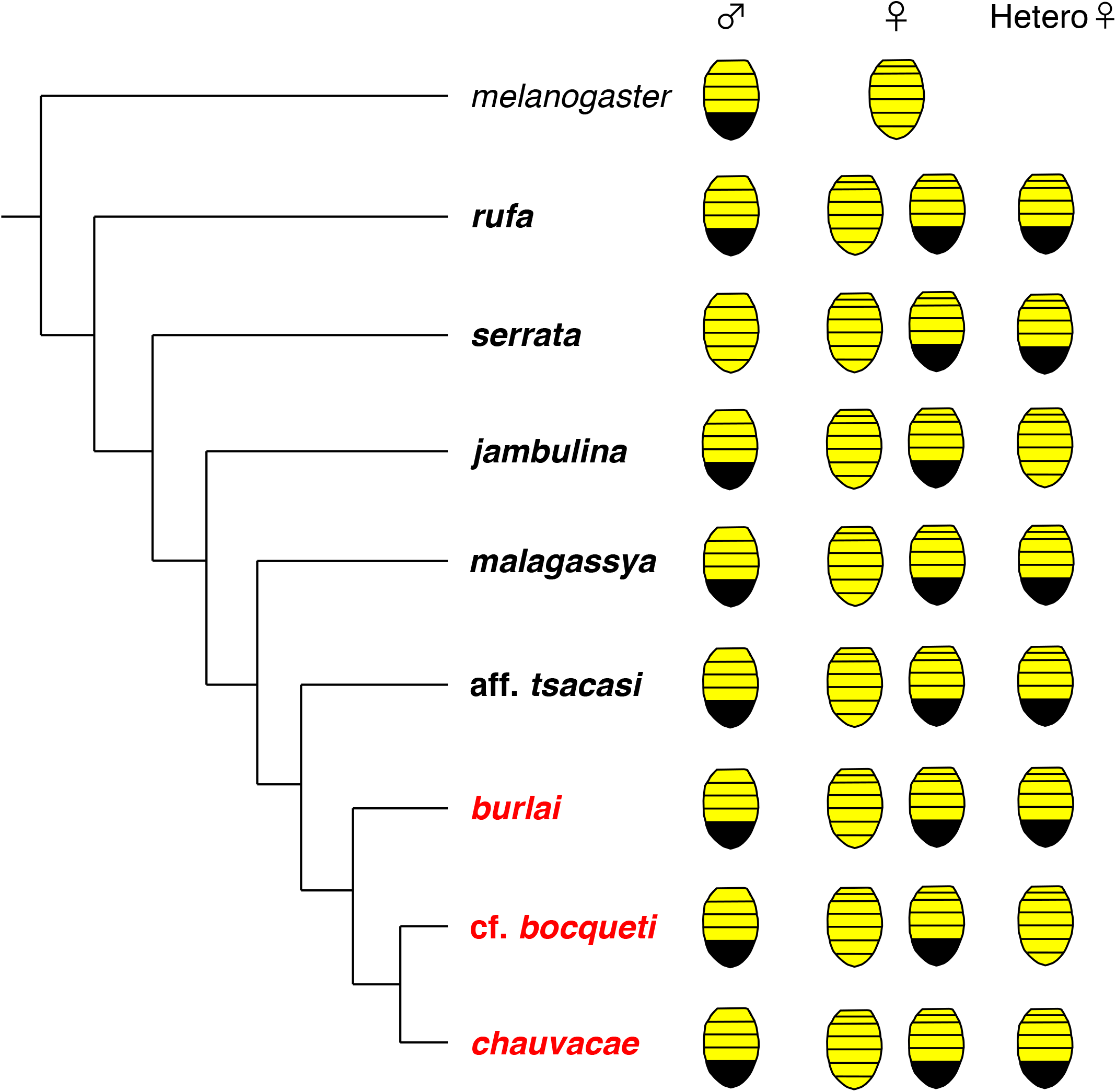
The phylogeny, phenotypes, and dominance relationships between abdominal color alleles in the *D. montium* species subgroup. The phylogeny is based on Conner et al. 2021; branch lengths are not to scale. Species in the *D. montium* clade are indicated in bold, and the species used in this study are highlighted in red. In each species, females show a single-locus Mendelian color dimorphism (Dark: the posterior abdominal segments are fully melanized; Light: the posterior segments show yellow pigmentation). In the rightmost column, the phenotype of female heterozygotes is indicated based on Yassin et al. 2016 and our crosses. The phenotype of *D. chauvacae* heterozygotes is based on personal communication from Dr. Amir Yassin, since no light strain of *D. chauvacae* is currently available in culture. In *D. jambulina* and *D.* cf. *bocqueti,* which are described as Light-dominant species, the Light *pdm3* allele is almost completely dominant at higher temperatures, and the Dark *pdm3* allele becomes increasingly dominant as the rearing temperature decreases (David et al 2026; Fukutomi et al. 2026).

We reasoned that if dominance is an intrinsic property of *pdm3* alleles, the same pair of alleles should retain the same dominance relationship when moved between genomic backgrounds. If, however, their dominance depends on the genotype at second-site modifiers, the same alleles will have different dominance in different backgrounds. We took advantage of the partial inter-fertility of some *montium* subgroup species to introgress Light *pdm3* alleles into the genomic backgrounds of Dark species or strains, and vice versa (Figure 2). These introgressions allowed us to compare females heterozygous for the same Light/Dark pair of *pdm3* alleles in homozygous Light, homozygous Dark, or heterozygous genomic backgrounds. Although not all potentially informative genotypes could be constructed due to hybrid sterility, this approach enabled us to test for the existence of second-site dominance modifiers in two pairs of closely related species: *D. burlai* vs *D. chauvacae*, and *D.* cf. *bocqueti* vs *D. chauvacae* (Table 1).

**Figure 2.**
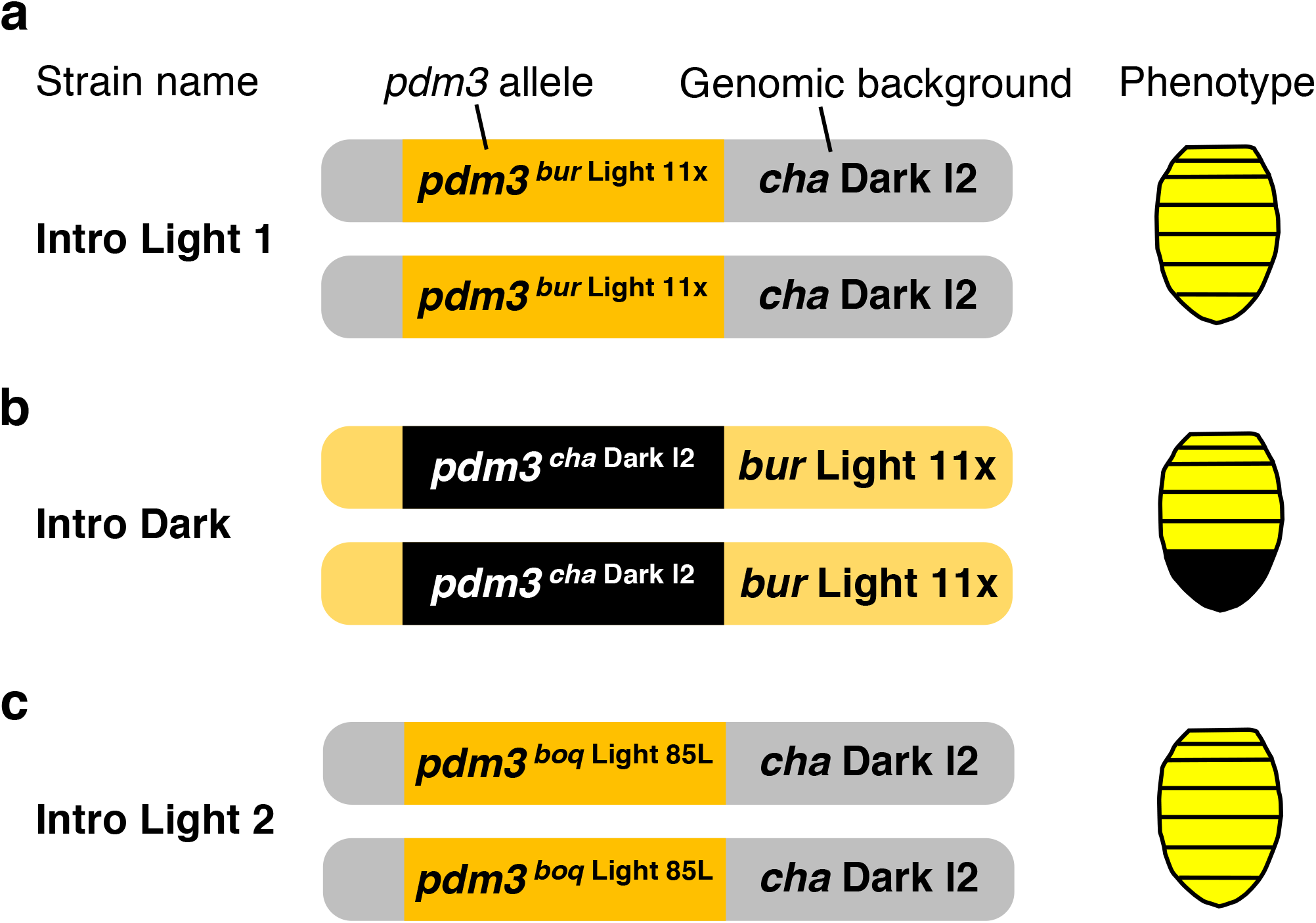
The genotypes and phenotypes of the introgression lines. **a,** “Intro Light 1” strain was created by introgressing the Light *pdm3* allele from *D. burlai* Light 11x strain into the genomic background of *D. chauvacae* Dark l2 strain. Flies from “Intro Light 1” strain show the light phenotype. **b,** “Intro Dark” strain was created by introgressing the Dark *pdm3* allele from *D. chauvacae* Dark l2 strain into the genomic background of *D. burlai* Light 11x strain. Flies from “Intro Dark” strain show the dark phenotype. **c,** “Intro Light 2” strain was created by introgressing the Light *pdm3* allele from *D. bocqueti* Light 85L strain into the genomic background of *D. chauvacae* Dark l2 strain. Flies from “Intro Light 2” strain show the light phenotype.

**Table 1.**
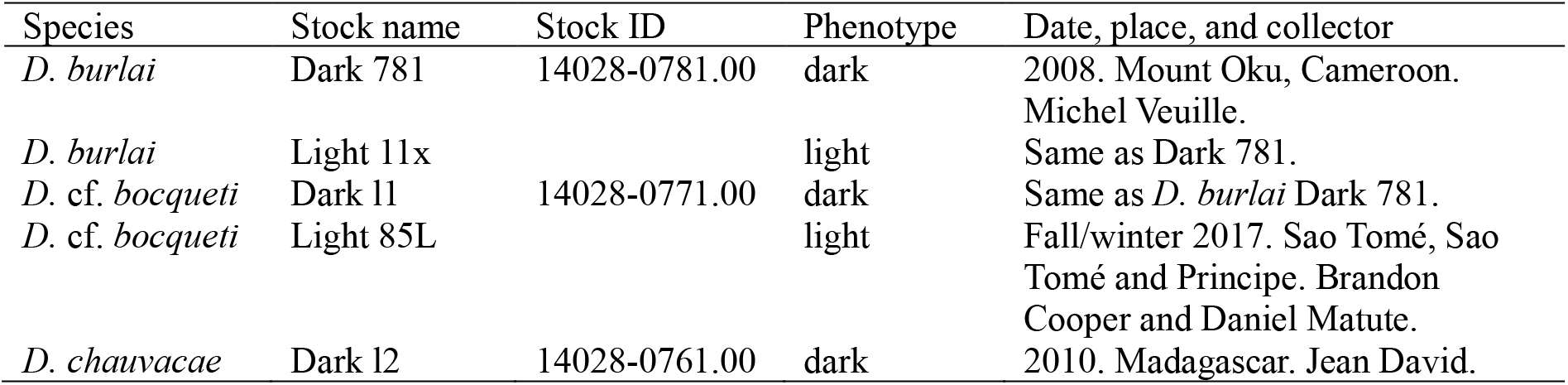
Strains used in this study. Each strain is homozygous for Dark or Light *pdm3* allele. Female phenotypes are indicated in the “Phenotype” column. For strains obtained from the US National Drosophila Species Stock Center, stock IDs are indicated.

## Results

### Dominance relationship between *D. chauvacae* and *D. burlai pdm3* alleles depends on the genomic background

In *D. burlai*, the Dark *pdm3* allele is completely dominant at all temperatures. However, the Light *pdm3* allele of *D. burlai* shows partial dominance at higher temperatures in the cross between *D. burlai* and *D. chauvacae* (Fukutomi et al. 2026; Fig. 3a). To test for the existence of dominance modifiers, we compared the A6 pigmentation of females heterozygous for the *D. burlai* Light and *D. chauvacae* Dark *pdm3* alleles in homozygous *burlai*-Light, homozygous *chauvacae*-Dark, and heterozygous genomic backgrounds. Phenotypic differences between the three genotypes were strong and significant (Figure 3; Table 2). The same Dark *pdm3* allele was more dominant over the same Light *pdm3* allele in the homozygous Dark genome (Fig. 3b) then in the homozygous Light genome (Fig. 3c), with the heterozygous genome showing an intermediate effect (Fig. 3a). While A6 pigmentation was temperature-dependent in the introgressions, as it is in the interspecific cross (Fukutomi et al. 2026), the genotype-by-temperature interaction was highly significant in every pairwise comparison (Table 2), confirming the effect of the genomic background. Significant effect of genomic background on the dominance relationship between the same pair of *pdm3* alleles was also observed in the A7 segment. As in the A6, the *burlai*-Light genome made the *D. burlai* Light *pdm3* allele more dominant, while the *chauvacae*-Dark genome made it less dominant (Figure S1); the difference in reaction norms was highly significant (Table S1). These results show that the dominance relationship between the *D. burlai* Light and *D. chauvacae* Dark *pdm3* alleles is strongly affected by second-site modifiers of dominance. Consistent with the species-specific plasticity of pigmentation in *pdm3* heterozygotes (Fukutomi et al. 2026; David et al. 2026), these dominance modifiers act in a temperature-dependent manner.

**Figure 3.**
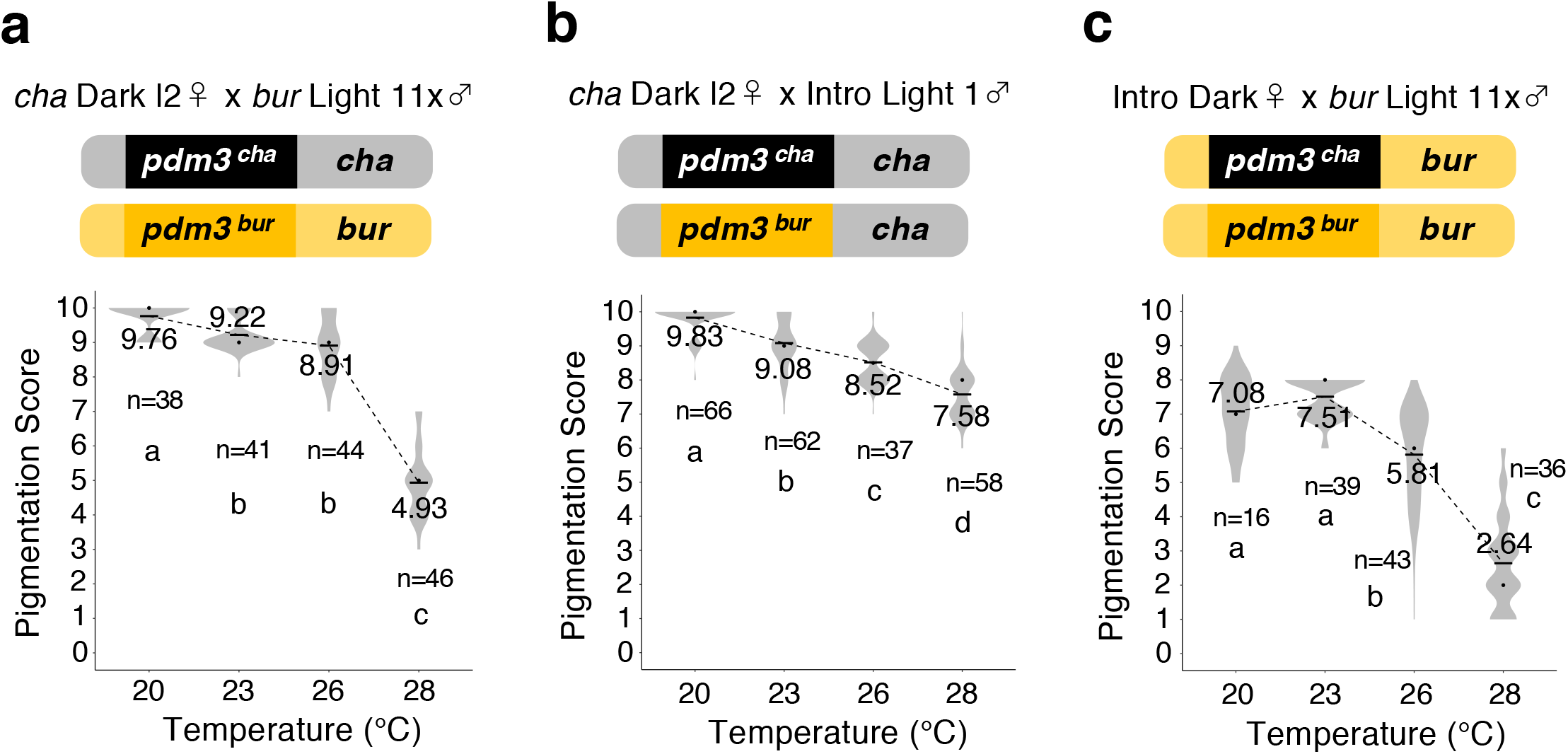
Pigmentation of *pdm3* heterozygotes depends on the genomic background of *D. chauvacae* or *D. burlai*. Female A6 pigmentation is shown for all genotypes. **a.** F1 females from the interspecific cross between females of the *D. chauvacae* Dark l2 strain and males of the *D. burlai* Light 11x strain. In this cross, both the *pdm3* locus and the genomic background are heterozygous. Significant differences between temperatures were detected by one-way ANOVA (degree of freedom = 3, *F* = 376.7, *p* < 10^-10^). **b.** F1 females from the cross between females of the *D. chauvacae* Dark l2 strain and males of the Intro Light 1 strain. Here, the *pdm3* locus is heterozygous, but the genomic background is mostly homozygous *D. chauvacae*. Significant differences between temperatures were detected by one-way ANOVA (degree of freedom = 3, *F* = 138.6, *p* < 10^-10^). **c.** F1 females from the cross between females of the Intro Dark strain and males of the *D. burlai* Light 11x strain. Here, the *pdm3* locus is heterozygous, but the genomic background is mostly homozygous *D. burlai*. Significant differences between temperatures were detected by one-way ANOVA (degree of freedom = 3, *F* = 217.9, *p* < 10^-10^). The reaction norms shown in **a**, **b**, and **c** are significantly different (two-way ANOVA, see Table 2).

**Table 2.**
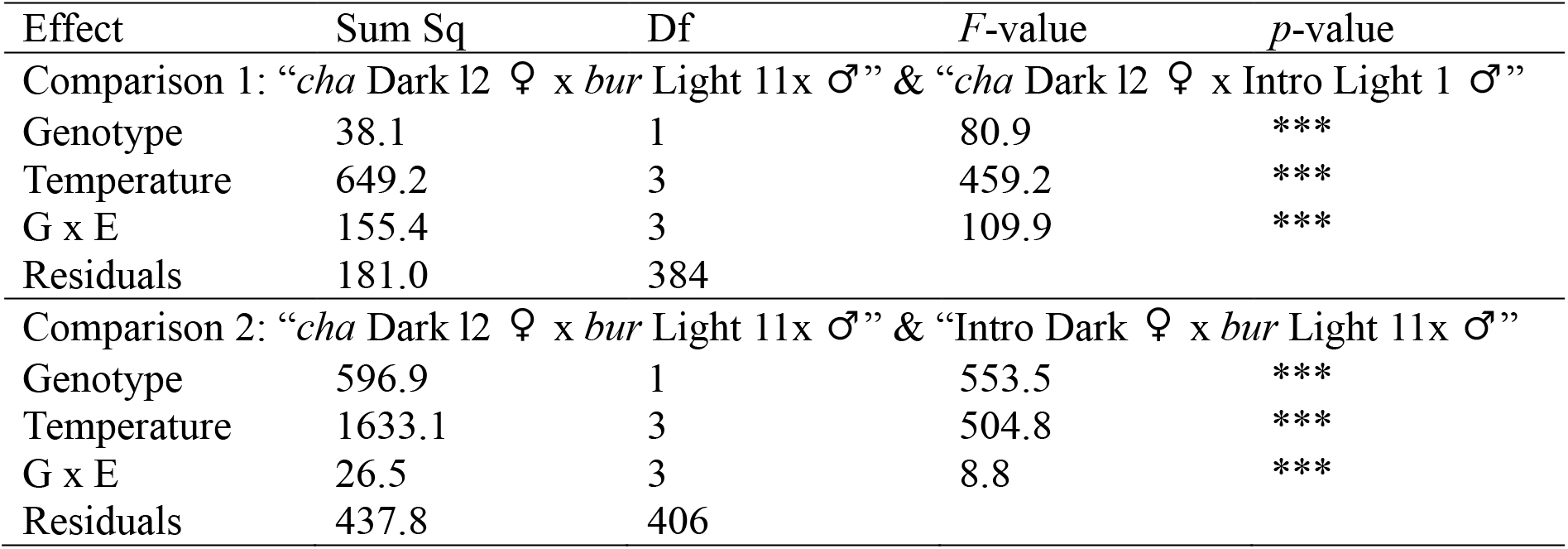
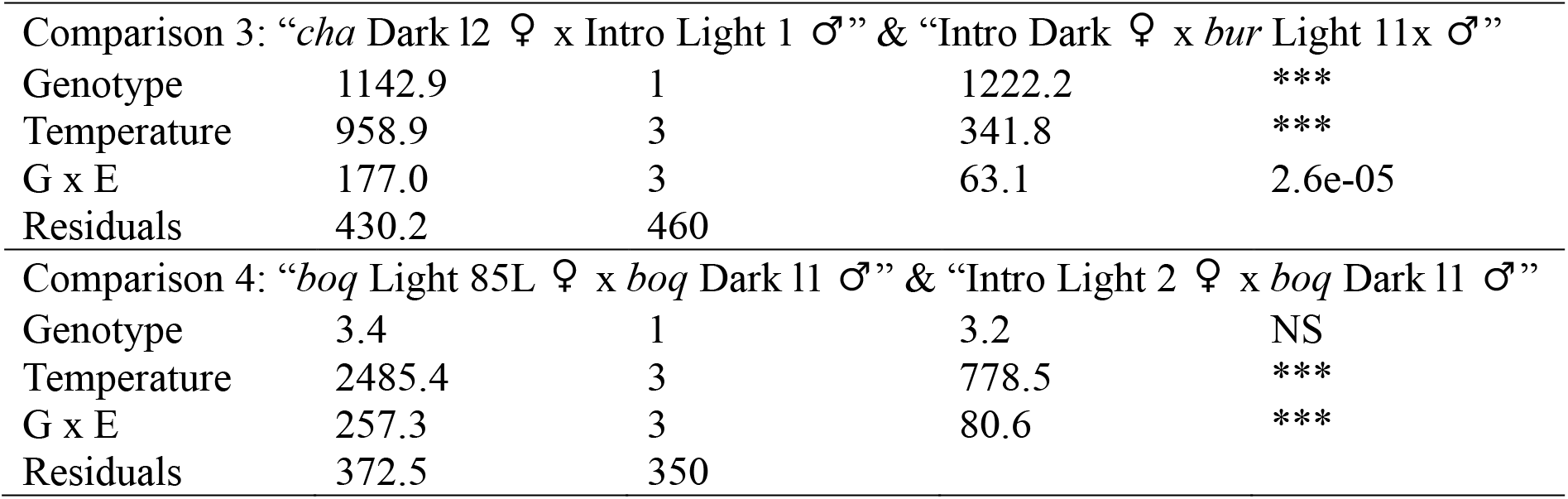
The results of two-way ANOVA analysis of A6 pigmentation in females heterozygous for Light and Dark *pdm3* alleles. Comparisons 1-3 contrast females carrying the *chauvacae* Dark *pdm3* allele and the *burlai* Light *pdm3* allele in different genomic backgrounds. For Comparison 4, females carrying the *bocqueti* Dark and Light *pdm3* alleles in different genomic backgrounds were analyzed. For the analysis in this comparison, 17 °C and 18 °C were treated as the same temperature. Five genotypes (shown in Figs. 3, 4) were used for the analyses: F1 females from the cross between *D. chauvacae* Dark l2 females and *D. burlai* Light 11x males, between *D. chauvacae* Dark l2 females and Intro Light 1 males, between Intro Dark females and *D. burlai* Light 11x males, between *D.* cf. *bocqueti* Light 85L females and *D.* cf. *bocqueti* Dark l1 males, and between Intro Light 2 females and *D.* cf. *bocqueti* Dark l1 males. Two “Genotypes” were compared in each analysis. “G x E” indicates an interaction between “Genotype” and “Temperature”. *cha*: *chauvacae*, *bur*: *burlai*, *boq*: *bocqueti*, Sum Sq: sum of squares, Df: degrees of freedom, ***: p <10^-10^, and NS: not significant.

In contrast to the modifiable dominance relationship between the *D. burlai* Light and *D. chauvacae* Dark *pdm3* alleles, the complete dominance of the *burlai* Dark *pdm3* allele over the *burlai* Light *pdm3* allele is insensitive to at least some changes in the genomic background. Completely dark A6 and A7 pigmentation is observed both in the *burlai*-Light / *burlai*-Dark heterozygous genome and in the *burlai*-Dark / *chauvacae*-Dark genome (Fig. S2; Table S2). This suggests that the effect of second-site dominance modifiers depends on the genotype at the main locus, with only some alleles being susceptible to modification.

### Dominance relationship between *D. chauvacae* and *D.* cf. *bocqueti pdm3* alleles depends on the genomic background

In *D.* cf. *bocqueti*, the Light *pdm3* allele is completely recessive at lower temperatures, and almost completely dominant at higher temperatures (Fukutomi et al. 2026). To test for the existence of dominance modifiers, we compared the A6 pigmentation of females heterozygous for the same *D.* cf. *bocqueti* Light and Dark *pdm3* alleles in different genomic backgrounds. We found that the dominance relationship between these *pdm3* alleles differed between the heterozygous *bocqueti-*Light / *bocqueti-*Dark genomic background and the *bocqueti-*Dark / *chauvacae*-Dark background. In the latter (all-Dark) genomic background, the Light *pdm3* allele was less dominant at the highest temperature, but more dominant at an intermediate temperature (Figure 4). A similar, temperature-dependent result was observed in the A7 segment (Fig. S3). In both segments, the genotype-by-temperature interaction was highly significant (Table 2; Table S3). These results demonstrate that, similar to the *D. burlai* / *D. chauvacae* species pair, second-site modifiers of dominance exist between *D.* cf. *bocqueti* and *D. chauvacae*.

**Figure 4.**
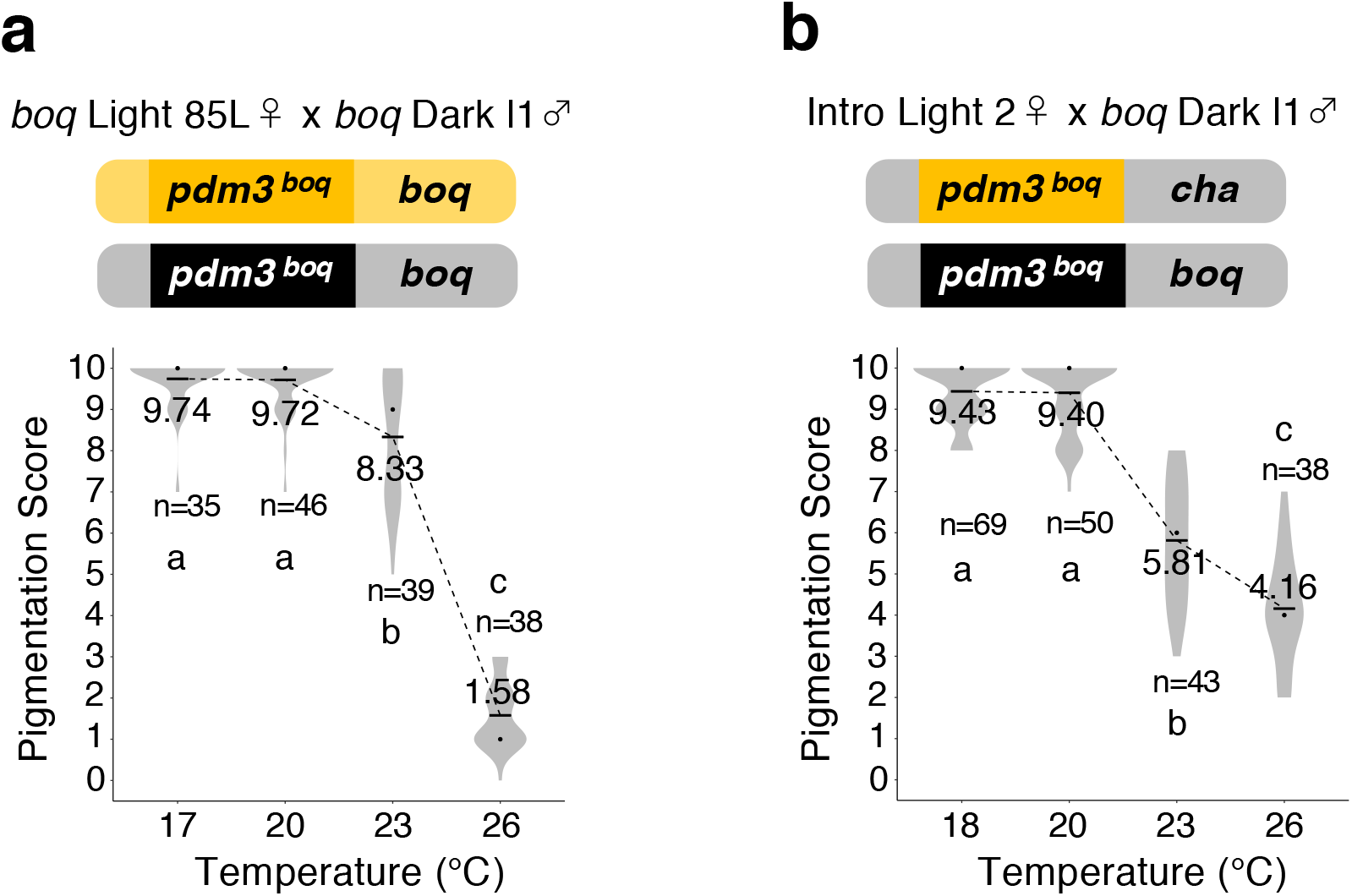
Pigmentation of *pdm3* heterozygotes depends on the genomic background of *D. chauvacae* or *D. bocqueti*. Female A6 pigmentation is shown for all genotypes. **a.** F1 females from the intraspecific cross between females of the *D. bocqueti* Light 85L strain and males of the *D. bocqueti* Dark l1 strain. Significant differences between temperatures were detected by one-way ANOVA (degree of freedom = 3, *F* = 636.7, *p* < 10^-10^). **b.** F1 females from the cross between females of the Intro Light 2 strain and males of the *D. bocqueti* Dark l1 strain. In this cross, the *pdm3* genotype is the same as in (a), but the genomic background is heterozygous *D. chauvacae* / *D. bocqueti*. Significant differences between temperatures were detected by one-way ANOVA (degree of freedom = 3, *F* = 278.1, *p* < 10^-10^). The reaction norms shown in **a** and **b** are significantly different (two-way ANOVA, see Table 2).

We then tested whether the genomic background of *D.* cf. *bocqueti* vs *D. chauvacae* affected the dominance relationship between the *D.* cf. *bocqueti* Light and the *D. chauvacae* Dark *pdm3* alleles. We observed significant phenotypic differences when this pair of alleles was placed in the homozygous *chauvacae*-Dark background, compared to the heterozygous *chauvacae*-Dark / *bocqueti*-Light background (Fig. S4; Table S4). In these genotypes, a key effect of the *chauvacae*-Dark genome was to increase the dominance of the *D.* cf. *bocqueti* Light *pdm3* allele at low temperature, while slightly decreasing its dominance at high temperature (Fig. S4). Together, these results show that the dominance effects of second-site modifiers depend both on the environmental conditions and, importantly, on the specific pair of alleles at the main locus controlling the phenotype.

## Conclusions

By holding the focal heterozygote constant while changing the rest of the genome, we show that dominance at a naturally polymorphic locus is shaped by interactions between focal alleles and their genomic background. In two species-pair comparisons, background substitution changed the phenotype of identical *pdm3* heterozygotes. These effects are not universal, but depended both on the particular pair of *pdm3* alleles and on environmental conditions. Dominance in this system is therefore a genotype-by-genotype-by-environment trait, rather than a fixed property of each allele. Both Light and Dark *pdm3* alleles segregate at high frequencies in natural populations (Ohnishi and Watanabe 1985; Yassin et al. 2016; Prigent et al. 2020), matching the key condition under which second-site dominance modifiers are expected to evolve (Bourguet 1999; Peischl and Bürger 2008; Llaurens et al 2015; Li and Bank 2024). Critically, our experiments show that these modifiers can alter dominance through their effects on phenotypic plasticity, providing a developmental mechanism for the evolution of dominance and broadening the range of conditions under which selection can favor the evolution of second-site modifiers.

The reciprocal introgression framework opens the way for understanding how dominance evolves across species. In the future, it will be important to identify the modifier loci, determine how derived modifier alleles affect dominance, reconstruct when these alleles arose relative to changes at *pdm3*, and examine how selection acted on these alleles. More broadly, these results link the evolution of dominance to epistasis and phenotypic plasticity (Li and Bank 2024), and suggest that functionally similar alleles may follow different evolutionary trajectories in different species.

## Materials and Methods

### *Drosophila* species and strains

*D. burlai*, *D. chauvacae*, and *D.* cf. *bocqueti* are closely related African species of the *D. montium* species subgroup (Yassin 2018, 2019; Conner et al., 2021). The strains used in this study are listed in Table 1. The males of all three species have dark abdominal pigmentation regardless of genotype; the Dark/Light dimorphism is only found in females (Figure 1) (Ohnishi and Watanabe 1985; Yassin et al 2016). Flies were reared on standard *Drosophila* cornmeal media, under a 12:12 hr day/night cycle, at temperatures from 17°C to 28°C. Not all species are viable at this full range of temperatures. To rear flies at 28°C, parental flies were crossed at 26°C and removed after two to three days; three day old or younger larvae were then moved to 28°C.

### Construction of introgression lines

The partial interfertility of *D. burlai*, *D. chauvacae*, and *D.* cf. *bocqueti* has allowed us to construct some, though not all, potentially informative introgression lines, where the Dark *pdm3* alleles were introgressed into the light genomic background and vice versa (Figure 2). All introgression crosses were conducted *en masse*. To make the *Intro Light 1* strain, the Light allele of *D. burlai* was introgressed into the genomic background of *D. chauvacae*, which has dark pigmentation. Females of the *D. chauvacae* Dark l2 strain were crossed to males of the *D. burlai* Light 11x strain, and F1 females were back-crossed to males of the *D. chauvacae* Dark l2 strain. In F2, light virgin females were selected and back-crossed again to *D. chauvacae* males. This procedure was repeated for eight generations. To make a pure-breeding light strain, subsequent generations were reared at 20°C, and light hybrid females were crossed to their sibling males for nine generations until all females showed the light phenotype. As the *burlai* Light *pdm3* allele is recessive to the *chauvacae* Dark allele at 20°C (Fukutomi et al. 2026), this allowed us to breed the *chauvacae pdm3* allele out of the introgression line despite the inability to detect that allele in males.

To make the *Intro Dark* strain, we introgressed the Dark *pdm3* allele of *D. chauvacae* into the genomic background of *D. burlai*. The first two generations of crosses were the same as for the Intro Light 1 strain. Subsequently, we selected dark virgin females, and back-crossed them to males of the *D. burlai* Light 11x strain. This procedure was repeated for 10 generations. To make a pure-breeding dark strain, we continued to select dark hybrid females and cross them to their sibling males for the following 15 generations, ensuring that all females showed the dark phenotype.

To make the *Intro Light 2* strain, the Light *pdm3* allele of *D*. cf. *bocqueti* was introgressed into the genomic background of *D. chauvacae*. Females of the *D. chauvacae* Dark l2 strain were crossed to males of the *D.* cf. *bocqueti* Light 85L strain, and F1 females were back-crossed to males of the *D. chauvacae* Dark l2 strain. In F2, light virgin females were selected and back-crossed to *D. chauvacae* males. This procedure was repeated for 15 generations. To make a pure-breeding light strain, subsequent generations were reared at 20°C, and light hybrid females were selected and crossed to their sibling males for another 15 generations. As the *bocqueti* Light allele is recessive to the *chauvacae* Dark allele at 20°C (Fukutomi et al. 2026), the phenotypically light females are homozygous with the Light *pdm3* allele. For the last several generations, we used introgression males homozygous for the *bocqueti* Light *pdm3* allele, which were identified by PCR on wing clips. Wings of individual males were placed directly in the PCR reaction solution, and a fragment of *pdm3* was amplified using primers GCTGAGAGAGGATTACGAGAGTC (forward) and TAATTAAGGTATGAGGAATAAGGCCTTTG (reverse). This pair of primers amplifies a 537 bp fragment from the *D. chauvacae*, but not from the *D.* cf. *bocqueti pdm3* allele (Fig. S5).

We note that introgression lines constructed in this manner provide a conservative test for second-site dominance modifiers because we cannot rule out that other genomic regions in addition to *pdm3* are introgressed across species. Thus, a positive result definitively shows the existence of second-site modifiers, while a negative result is inconclusive because we cannot rule out the existence of modifiers that are closely linked to the *pdm3* locus.

### Confirming the genetic association in *D. chauvacae*

The female-limited, single-gene Mendelian color polymorphism in the *montium* subgroup is associated with *pdm3* in all tested species including *D. burlai* and *D.* cf. *bocqueti* (Yassin et al 2016; Fukutomi et al. 2026). To test whether this polymorphism is also linked to the *pdm3* locus in *D. chauvacae*, we genotyped dark and light females from advanced introgression strains between *D. chauvacae* and *D. burlai,* and between *D. chauvacae* and *D.* cf. *bocqueti*.

For the *D. chauvacae* / *D. burlai* genotyping, we first identified interspecific differences by sequencing a 439 bp region around Exon 5 of *pdm3*. The region was amplified with primers TCGGAGCAGTTCTTGCCCAATG (forward) and GATCGGCAGCCAGATGCGAG (reverse) from four females of the *chauvacae* Dark l2 strain and four females of the *burlai* Light 11x strain. The restriction enzyme EarI (New England Biolabs) cuts the *burlai* sequence, but not the *chauvacae* sequence (Fig. S6a). We used PCR followed by EarI digest to genotype 22 dark and 19 light females from the Intro Dark strain, where the Dark allele of *D. chauvacae* was introgressed into the genomic background of *D. burlai* (see previous section). All dark females were homozygous for the *chauvacae pdm3* allele, while all light females carried the *burlai* Light 11x allele (Fig. S6b-e).

The *D. chauvacae* / *D.* cf. *bocqueti* introgression was genotyped by sequencing. To identify differences between parental strains, we amplified and sequenced a 770 bp region containing Exon 11 of *pdm3* from the *D. chauvacae* Dark l2 strain and the *bocqueti* Light 85L strain using primers ACCGAGAGAGAGCCGCGTTTC (forward) and CTCGGGTTCTGTGCTTCGCTTG (reverse) (Fukutomi et al. 2026). Subsequently, the same region was amplified and sequenced from 20 light and 21 dark females from the Intro Light 2 strain, where the Light allele of *D.* cf. *bocqueti* was introgressed into the genomic background of *D. chauvacae* (see previous section). The sequences were aligned with MEGA11 (Tamura et al. 2021). All light introgression females had SNP alleles specific to the *bocqueti* Light 85L strain, and all dark females had alleles specific to the *chauvacae* Dark l2 strain (Table S5).

### Quantifying abdominal pigmentation

Females heterozygous for *pdm3* were obtained from mass crosses (30 parental females * 30 parental males), aged for 6-8 days post-eclosion, and fixed in 100 % ethanol. The pigmentation of the 6^th^ and 7^th^ abdominal segments (A6 and A7) of adult females was scored on the scale from 0 (completely yellow) to 10 (completely black) as described in David et al. 1990. The scoring was conducted blindly, so that the scorers did not know the genotypes or the rearing temperatures. The graphs of pigmentation scores were generated by ggplot2 (Wickham 2009). Statistical analyses were conducted with R version 4.2.1 (R Core Team 2022). To test whether the scores were significantly different between rearing temperatures in the same cross, one-way ANOVA and Tukey’s honest significant differences (HSD) were used. To compare reaction norms between different genotypes, we used two-way ANOVA. In two-way ANOVA analysis, the following data published in Fukutomi et al. 2026 were reused; Data of 20°C, 23°C, and 26°C in Fig. 3a, Fig. S1a, Fig. S2a, c, Fig. S4a, c, all the temperature data in Fig. 4a, and in Fig. S3a.

## Supplementary Tables

**Table S1.** The results of two-way ANOVA of A7 pigmentation in females heterozygous for the *chauvacae* Dark *pdm3* allele and the *burlai* Light *pdm3* allele in different genomic backgrounds. Three genotypes (Fig. S1) were used for the analyses: F1 females from the cross between *D. chauvacae* Dark l2 females and *D. burlai* Light 11x males, between *D. chauvacae* Dark l2 females and Intro Light 1 males, between Intro Dark females and *D. burlai* Light 11x males. Two “Genotypes” were compared in each analysis. “G x E” indicates an interaction between “Genotype” and “Temperature”. *cha*: *chauvacae*, *bur*: *burlai*, Sum Sq: sum of squares, Df: degrees of freedom, ***: *p* <10^-10^.

| Effect | Sum Sq | Df | F-value | p-value |
| --- | --- | --- | --- | --- |
| Comparison 1: “*cha* Dark l2 ♀ x *bur* Light 11x ♂” & “*cha* Dark l2 ♀ x Intro Light 1 ♂” |  |  |  |  |
| Genotype | 118.0 | 1 | 77.6 | *** |
| Temperature | 1744.3 | 3 | 382.2 | *** |
| G x E | 52.4 | 3 | 11.5 | 3.2e-07 |
| Residuals | 584.2 | 384 |  |  |
| Comparison 2: “*cha* Dark l2 ♀ x *bur* Light 11x ♂” & “Intro Dark ♀ x *bur* Light 11x ♂” |  |  |  |  |
| Genotype | 126.6 | 1 | 97.6 | *** |
| Temperature | 1248.8 | 3 | 320.9 | *** |
| G x E | 161.6 | 3 | 41.5 | *** |
| Residuals | 526.6 | 406 |  |  |
| Comparison 3: “*cha* Dark l2 ♀ x Intro Light 1 ♂” & “Intro Dark ♀ x *bur* Light 11x ♂” |  |  |  |  |
| Genotype | 599.7 | 1 | 529.5 | *** |
| Temperature | 1243.6 | 3 | 366.0 | *** |
| G x E | 121.4 | 3 | 35.7 | *** |
| Residuals | 521.0 | 460 |  |  |

**Table S2.** The results of two-way ANOVA of A6 and A7 pigmentation in females heterozygous for the *burlai* Dark and Light *pdm3* alleles in different genomic backgrounds (Fig. S2). Comparison 1 is for A6 pigmentation and Comparison 2 is for A7 pigmentation. F1 females from the cross between *D. burlai* Light 11x females and *D. burlai* Dark 781 males, and F1 females from the cross between Intro Light 1 females and *D. burlai* Dark 781 males, were contrasted in each comparison. “G x E” indicates an interaction between “Genotype” and “Temperature”. Sum Sq: sum of squares, Df: degrees of freedom, NS: not significant.

| Effect | Sum Sq | Df | F-value | p-value |
| --- | --- | --- | --- | --- |
| Comparison 1: A6 pigmentation |  |  |  |  |
| Genotype | 0.012 | 1 | 2.2 | NS |
| Temperature | 0.015 | 2 | 1.4 | NS |
| G x E | 0.016 | 2 | 1.6 | NS |
| Residuals | 1.947 | 380 |  |  |
| Comparison 2: A7 pigmentation |  |  |  |  |
| Genotype | 0.66 | 1 | 3.1 | 0.048 |
| Temperature | 0.23 | 2 | 17.6 | 3.4e-05 |
| G x E | 0.22 | 2 | 2.9 | NS |
| Residuals | 14.23 | 380 |  |  |

**Table S3.** The results of two-way ANOVA of A7 pigmentation in females heterozygous for the *D. bocqueti* Dark and Light *pdm3* alleles in different genomic backgrounds. Two genotypes (shown in Fig. S3) were compared: F1 females from the cross between *D.* cf. *bocqueti* Light 85L females and *D.* cf. *bocqueti* Dark l1 males, and between Intro Light females and *D.* cf. *bocqueti* Dark l1 males. For this analysis, 17 °C and 18 °C were treated as the same temperature. “G x E” indicates an interaction between “Genotype” and “Temperature”. Sum Sq: sum of squares, Df: degrees of freedom, ***: *p* <10^-10^.

| Effect | Sum Sq | Df | F-value | p-value |
| --- | --- | --- | --- | --- |
| Genotype | 43.4 | 1 | 10.0 | *** |
| Temperature | 1623.6 | 3 | 124.6 | 0.0017 |
| G x E | 236.2 | 3 | 18.2 | *** |
| Residuals | 1520.7 | 350 |  |  |

**Table S4.** The results of two-way ANOVA of A6 and A7 pigmentation in females heterozygous for the *chauvacae* Dark and the *bocqueti* Light *pdm3* alleles in different genomic backgrounds (Fig. S4). Comparison 1 is for A6 pigmentation and Comparison 2 is for A7 pigmentation. Each comparison contrasts F1 females from the cross between *D. chauvacae* Dark l2 females and *D.*cf. *bocqueti* Light 85L males, and F1 females from the cross between *D. chauvacae* Dark l2 females and Intro Light 2 males. “G x E” indicates an interaction between “Genotype” and “Temperature”. Sum Sq: sum of squares, Df: degrees of freedom, NS: not significant.

| Effect | Sum Sq | Df | F-value | p-value |
| --- | --- | --- | --- | --- |
| Comparison 1: A6 pigmentation |  |  |  |  |
| Genotype | 5.0 | 1 | 3.0 | NS |
| Temperature | 1977.1 | 2 | 593.0 | *** |
| G x E | 105.0 | 2 | 31.5 | *** |
| Residuals | 508.4 | 305 |  |  |
| Comparison 2: A7 pigmentation |  |  |  |  |
| Genotype | 15.0 | 1 | 12.6 | 0.00045 |
| Temperature | 1738.5 | 2 | 731.1 | *** |
| G x E | 67.5 | 2 | 28.4 | *** |
| Residuals | 362.7 | 305 |  |  |

**Table S5.** The results of genotyping in the Intro Light 2 strain, where the Light allele of *D.* cf. *bocqueti* was introgressed into the genomic background of *D. chauvacae*. A 770 bp PCR amplicon around Exon 11 of *pdm3* contains five single nucleotide polymorphisms (SNPs) which distinguish the light paternal strain *bocqueti* Light 85L and the dark parental strain *chauvacae* Dark l2. SNP alleles and positions relative to the 5’ end of the amplicon are indicated in the “Position” columns. All light females from the introgression line (Light Intro females) had the same sequence as the Light 85L parental strain. All dark females from the introgression line (Dark Intro females) had the SNPs specific to the Dark l2 parental strain, as well as three SNPs which did not appear in either parental strain. The latter three SNPs may exist in Dark l2 strain at low frequency, leading us to miss them in the four parental individuals.

| Individuals | Position |  |  |  |  |  |  |  |
| --- | --- | --- | --- | --- | --- | --- | --- | --- |
|  | 190 | 201 | 203 | 212 | 276 | 282 | 357 | 441 |
| *Light 85L* (n=4) | C | C | G | C | T | C | C | C |
| *Dark l2* (n=4) | T | T | T | C | C | C | C | T |
| Light Intro females (n=20) | C | C | G | C | T | C | C | C |
| Dark Intro females (n=21) | T | T | T | A | C | C (n=12) / G (n=9) | T | T |

## Supplementary figure legends

**Figure S1**

Pigmentation of *pdm3* heterozygotes depends on the genomic background of *D. chauvacae* or *D. burlai*. Female A7 pigmentation is shown for all genotypes. **a.** F1 females from the interspecific cross between females of the *D. chauvacae* Dark l2 strain and males of the *D. burlai* Light 11x strain. In this cross, both the *pdm3* locus and the genomic background are heterozygous. Significant differences between temperatures were detected by one-way ANOVA (degree of freedom = 3, *F* = 175.6, *p* < 10^-10^). **b.** F1 females from the cross between females of the *D. chauvacae* Dark l2 strain and males of the Intro Light 1 strain. Here, the *pdm3* locus is heterozygous, but the genomic background is mostly homozygous *D. chauvacae*. Significant differences between temperatures were detected by one-way ANOVA (degree of freedom = 3, *F* = 216.5, *p* < 10^-10^). **c.** F1 females from the cross between females of the Intro Dark strain and males of the *D. burlai* Light 11x strain. Here, the *pdm3* locus is heterozygous, but the genomic background is mostly homozygous *D. burlai*. Significant differences between temperatures were detected by one-way ANOVA (degree of freedom = 3, *F* = 144.3, *p* < 10^-10^). The reaction norms shown in **a**, **b**, and **c** are significantly different (two-way ANOVA, see Table S1).

**Figure S2**

Pigmentation of females heterozygous for the Dark and Light *pdm3* alleles of *D*. *burlai* does not depend on the genomic background of *D. chauvacae* or *D. burlai*. **a.** A6 pigmentation of F1 females from the intraspecific cross between females of the *D. burlai* Light 11x strain and males of the *D. burlai* Dark 781 strain. In this cross, the genomic background is homozygous *D. burlai*. No significant difference between temperatures was detected by one-way ANOVA (degree of freedom = 2, *F* = 1.05). **b.** A6 pigmentation of F1 females from the cross between females of the Intro Light 1 strain and males of the *D. burlai* Dark 781 strain. In this cross, the genomic background is mostly heterozygous *D. chauvacae* / *D. burlai*. No significant difference between temperatures was detected by one-way ANOVA (degree of freedom = 2, *F* = 1.61). **c.** A7 pigmentation of F1 females from the intraspecific cross between females of the *D. burlai* Light 11x strain and males of the *D. burlai* Dark 781 strain. In this cross, the genomic background is homozygous *D. burlai*. No significant difference between temperatures was detected by one-way ANOVA (degree of freedom = 2, *F* = 1.05). **d.** A7 pigmentation of F1 females from the cross between females of the Intro Light 1 strain and males of the *D. burlai* Dark 781 strain. In this cross, genomic background is mostly heterozygous *D. chauvacae* / *D. burlai*. A slight difference between temperatures was detected by one-way ANOVA (degree of freedom = 2, *F* = 3.21, *p* = 0.043). The comparisons between reaction norms in **a** and **b,** and between reaction norms in **c** and **d,** did not show significant differences (two-way ANOVA, see Table S2).

**Figure S3**

Pigmentation of *pdm3* heterozygotes depends on the genomic background of *D. chauvacae* or *D.* cf*. bocqueti*. Female A7 pigmentation is shown for all genotypes. **a.** F1 females from the intraspecific cross between females of *D.* cf. *bocqueti* Light 85L strain and males of *D.* cf. *bocqueti* Dark l1 strain. In this cross, the genomic background is homozygous *D.* cf. *bocqueti*. Significant difference between temperatures was detected by one-way ANOVA (degree of freedom = 3, *F* = 42.9, *p* = 10^-10^). **b.** F1 females from the cross between females of Intro Light 2 strain and males of Dark l1 strain. In this cross, the genomic background is mostly heterozygous *D. chauvacae* / *D.* cf. *bocqueti*. Significant difference between temperatures was detected by one-way ANOVA (degree of freedom = 3, *F* = 289.7, *p* < 10 ^-10^). The reaction norms shown in a and b are significantly different (two-way ANOVA, see Table S3).

**Figure S4**

Pigmentation of females heterozygous for the *chauvacae* Dark and the *bocqueti* Light *pdm3* alleles depends on the genomic background of *D. chauvacae* or *D.* cf. *bocqueti*. **a.** A6 pigmentation of F1 females from the cross between females of the *D. chauvacae* Dark l2 strain and males of *D.* cf. *bocqueti* Light 85L strain. In this cross, the genomic background is heterozygous *D. chauvacae* / *D.* cf. *bocqueti*. Significant difference between temperatures was detected by one-way ANOVA (degree of freedom = 2, *F* = 405.4, *p* < 10^-10^). **b.** A6 pigmentation of F1 females from the cross between females of the Dark l2 strain and males of the Intro Light 2 strain. In this cross, the genomic background is mostly homozygous *D. chauvacae*. Significant difference between temperatures was detected by one-way ANOVA (degree of freedom = 2, *F* = 196.0, *p* < 10^-10^). **c.** A7 pigmentation of F1 females from the cross between females of the *D. chauvacae* Dark l2 strain and males of *D.* cf. *bocqueti* Light 85L strain. In this cross, the genomic background is heterozygous *D. chauvacae* / *D.* cf. *bocqueti*. Significant difference between temperatures was detected by one-way ANOVA (degree of freedom = 2, *F* = 390.3, *p* < 10^-10^). **d.** A7 pigmentation of F1 females from the cross between females of the Dark l2 strain and males of the Intro Light 2 strain. In this cross, the genomic background is mostly homozygous *D. chauvacae*. Significant difference between temperatures was detected by one-way ANOVA (degree of freedom = 2, *F* = 289.7, *p* < 10^-10^).

**Figure S5**

PCR distinguishes the *pdm3* alleles of *D. chauvacae* and *D.* cf. *bocqueti*. The results of PCR on male wings dissected from *D. chauvacae* and *D.* cf. *bocqueti*. The numbers indicate lanes on the agarose gel. The samples in each lane are as follows. Lane 1: GeneRuler 100 bp DNA Ladder (Thermo Fisher Scientific). Lanes 2-3: PCR products amplified from two different males of the *D. chauvacae* Dark l2 strain. Lanes 4-5: PCR products amplified from two different males of the *D.* cf. *bocqueti* Light 85L strain.

**Figure S6**

Restriction enzyme EarI distinguishes the *pdm3* alleles of *D. burlai* and *D. chauvacae*. **a.** EarI digests of PCR amplicons of *D. burlai*, *D. chauvacae*, and their F1 hybrids. Lane 1: O’RangeRuler 20bp DNA Ladder (Thermo Fisher Scientific). Lane 2: PCR amplicon from *D. burlai* Light 11x females (undigested). Lane 3: PCR amplicon from *D. burlai* Light 11x females digested with EarI. Lane 4: PCR amplicon from *D. chauvacae* Dark l2 females (undigested). Lane 5: PCR amplicon from *D. chauvacae* Dark l2 females digested with EarI. Lane 6: PCR amplicon from F1 females from the cross between Dark l2 females and Light 11x males (undigested). Lane 7: PCR amplicon from F1 females from the cross between Dark l2 females and Light 11x males, digested with EarI. **b**. Genotyping of light females from the Intro Dark strain. Lane 1: O’RangeRuler 20bp DNA Ladder (Thermo Fisher Scientific). Lane 2-11: PCR amplicons digested with EarI. **c**. Additional light females (lanes 3-11). Lane 1: 1kb Plus DNA Ladder (New England Biolabs). Lane 2: O’RangeRuler 20bp DNA Ladder (Thermo Fisher Scientific). **d.** Genotyping of dark females from the Intro Dark strain. Lane 1: 1kb Plus DNA Ladder (New England Biolabs). Lane 2-12: PCR amplicons digested with EarI. **e**. Additional dark females (lanes 2-12). Lane 1: 1kb Plus DNA Ladder (New England Biolabs).

## Data, Materials, and Software Availability

Raw phenotypic data and R scripts will be uploaded to Figshare (https://figshare.com/). Sequencing data will be deposited to DDBJ (ddbj.nig.ac.jp).

## Acknowledgements

We thank Jean R. David, Brandon Cooper, Daniel Matute, and the US National *Drosophila* Species Stock Center for providing *Drosophila* strains; David Begun, Susan Lott, and Joanna Chiu for the use of their incubators; Amir Yassin for advice on taxonomy; and Michael Turelli for comments on the manuscript. This work was funded by NIH grant R35 GM122592 to AK and Research Fellowship for Young Scientists KAKENHI 21J00655 from the Japan Society for the Promotion of Science to YF.

## Author contributions

Y.F. and A.K. designed the research; Y.F., A.P.G., and J.Z. collected data; Y.F. analyzed data; Y.F. and A.K. wrote the paper.

## References

Barson NJ, Aykanat T, Hindar K, Baranski M, Bolstad GH, Fiske P, Jacq C, Jensen AJ, Johnston SE, Karlsson S, et al. 2015. Sex-dependent dominance at a single locus maintains variation in age at maturity in salmon. Nature. 528(7582):405–408. doi:10.1038/nature16062.

Belt AL, Burnet B. 1972. Experimental modification of the dominance relations of a melanotic tumour gene in Drosophila melanogaster. Genet Res. 20(2):115–132. doi:10.1017/S0016672300013628.

Billiard S, Castric V. 2011. Evidence for Fisher’s dominance theory: how many ‘special cases’? Trends Genet. 27(11):441–445. doi:10.1016/j.tig.2011.06.005.

Billiard S, Castric V, Llaurens V. 2021. The integrative biology of genetic dominance. Biol Rev. 96(6):2925–2942. doi:10.1111/brv.12786.

Bourguet D. 1999. The evolution of dominance. Heredity. 83:1–4. doi:10.1038/sj.hdy.6885600.

Bourguet D, Prout M, Raymond M. 1996. Dominance of Insecticide Resistance Presents a Plastic Response. Genetics. 143, 407–416. doi:10.1093/genetics/143.1.407.

Conner WR, Delaney EK, Bronski MJ, Ginsberg PS, Wheeler TB, Richardson KM, Peckenpaugh B, Kim KJ, Watada M, Hoffmann AA, et al. 2021. A phylogeny for the Drosophila montium species group: a model clade for comparative analyses. Mol Phylogenet Evol. 158:107061. doi:10.1016/j.ympev.2020.107061.

David JR, Capy P, Gauthier JP. 1990. Abdominal pigmentation and growth temperature in Drosophila melanogaster: similarities and differences in the norms of reaction of successive segments. J Evol Biol. 3(5–6):429–445. doi:10.1046/j.1420-9101.1990.3050429.x.

David JR, Denis B, Francelle P, Lemaire A, Das A, Mohanty S, Gibert P, Yassin A. 2026. Phenotypic plasticity as a function of genetic polymorphism: thermal dominance reversal in Drosophila species with contrasting melanism. Evolution. 80(1):127–142. doi:10.1093/evolut/qpaf225.

Di C, Lohmueller KE. 2024. Revisiting dominance in population genetics. Genome Biol Evol. 16(8):evae147. doi:10.1093/gbe/evae147.

Durand E, Méheust R, Soucaze M, Goubet PM, Gallina S, Poux C, Fobis-Loisy I, Guillon E, Gaude T, Sarazin A, et al. 2014. Dominance hierarchy arising from the evolution of a complex small RNA regulatory network. Science. 346(6214):1200–1205. doi:10.1126/science.1259442.

Fisher RA. 1928. The possible modification of the response of the wild type to recurrent mutations. Am Nat. 62(679):115–126. doi:10.1086/280193.

Fukutomi Y, Phillips-Garcia A, Liu J, Chuang A, Watada M, Ramniwas S, Kopp A. 2026. Evolution of dominance in a Mendelian trait is linked to the evolution of environmental plasticity. Evolution. 80(1):108–126. doi:10.1093/evolut/qpaf185.

Green RM, Fish JL, Young NM, Smith FJ, Roberts B, Dolan K, Choi I, Leach CL, Gordon P, Cheverud JM, et al. 2017. Developmental nonlinearity drives phenotypic robustness. Nat Commun. 8:1970. doi:10.1038/s41467-017-02037-7

Grieshop K, Arnqvist G. 2018. Sex-specific dominance reversal of genetic variation for fitness. PLoS Biol. 16(12):e2006810. doi:10.1371/journal.pbio.2006810.

Grieshop K, Ho EKH, Kasimatis KR. 2024. Dominance reversals: the resolution of genetic conflict and maintenance of genetic variation. Proc R Soc B. 291(2018):20232816. doi:10.1098/rspb.2023.2816.

Groh JS, Ackerman G, Wenzel M, Coop G. 2025. Distinct haplotypes and reversed dominance at a single-gene balanced polymorphism controlling heterodichogamous flowering in two genera of wingnuts. Curr Biol. 35(21):5192–5209.e11. doi:10.1016/j.cub.2025.09.061.

Guio L, González J. 2015. The dominance effect of the adaptive transposable element insertion Bari-Jheh depends on the genetic background. Genome Biol Evol. 7(5):1260–1266. doi:10.1093/gbe/evv071.

Haldane JBS. 1930. A note on Fisher’s theory of the origin of dominance, and on a correlation between dominance and linkage. Am Nat. 64(690):87–90. doi:10.1086/280299.

Karageorgi M, Lyulina AS, Bitter MC, et al. 2025. Beneficial reversal of dominance maintains a large-effect resistance polymorphism under fluctuating insecticide selection. Nat Ecol Evol. 9:2145–2160. doi:10.1038/s41559-025-02853-x.

Klingenberg CP. 2019. Phenotypic plasticity, developmental instability, and robustness: the concepts and how they are connected. Front Ecol Evol. 7:56. doi:10.3389/fevo.2019.00056.

Le Poul Y, Whibley A, Chouteau M, Prunier F, Llaurens V, Joron M. 2014. Evolution of dominance mechanisms at a butterfly mimicry supergene. Nat Commun. 5:5644. doi:10.1038/ncomms6644.

Li J, Bank C. 2024. Dominance and multi-locus interaction. Trends Genet. 40(4):364–378. doi:10.1016/j.tig.2023.12.003.

Llaurens V, Joron M, Billiard S. 2015. Molecular mechanisms of dominance evolution in Müllerian mimicry. Evolution. 69(12):3097–3108. doi:10.1111/evo.12810.

Mikkola K. 1984. Dominance relations among the melanic forms of Biston betularius and Odontopera bidentata (Lepidoptera, Geometridae). Heredity. 52:9–16. doi:10.1038/hdy.1984.2.

Naisbit RE, Jiggins CD, Mallet J. 2003. Mimicry: developmental genes that contribute to speciation. Evol Dev. 5(3):269–280. doi:10.1046/j.1525-142X.2003.03034.x.

Nijhout HF. 2003. Polymorphic mimicry in Papilio dardanus: mosaic dominance, big effects, and origins. Evol Dev. 5(6):579–592. doi:10.1046/j.1525-142X.2003.03063.x.

Ohnishi S, Watanabe TK. 1985. Genetic analysis of color dimorphism in the Drosophila montium subgroup. Jpn J Genet. 60(4):355–358. doi:10.1266/jjg.60.355.

Pearse DE, Barson NJ, Nome T, Gao G, Campbell MA, Abadía-Cardoso A, Anderson EC, Rundio DE, Williams TH, Naish KA, et al. 2019. Sex-dependent dominance maintains migration supergene in rainbow trout. Nat Ecol Evol. 3:1731–1742. doi:10.1038/s41559-019-1044-6.

Peischl S, Bürger R. 2008. Evolution of dominance under frequency-dependent intraspecific competition. J Theor Biol. 251(2):210–226. doi:10.1016/j.jtbi.2007.11.014.

Phadke SS, Paixão T, Pham T, Pham S, Zufall RA. 2014. Genetic background alters dominance relationships between mat alleles in the ciliate Tetrahymena thermophila. J Hered. 105(1):130–135. doi:10.1093/jhered/est063.

Posavi M, Gelembiuk GW, Larget B, Lee CE. 2014. Testing for beneficial reversal of dominance during salinity shifts in the invasive copepod Eurytemora affinis, and implications for the maintenance of genetic variation. Evolution. 68(11):3166–3183. doi:10.1111/evo.12502.

Prigent SR, Lang M, Nagy O, Acurio A, Matamoro-Vidal A, Courtier-Orgogozo V, David JR. 2020. Field collections reveal that São Tomé is the Afrotropical island with the highest diversity of drosophilid flies (Diptera: Drosophilidae). Ann Soc Entomol Fr (N.S.). 56(1):1–14. doi:10.1080/00379271.2019.1703814.

R Core Team. 2022. R: A language and environment for statistical computing. Vienna (Austria): R Foundation for Statistical Computing. https://www.R-project.org/.

Rendel JM. 1959. Canalization of the scute phenotype of Drosophila. Evolution. 13(4):425–439. doi:10.1111/j.1558-5646.1959.tb03033.x.

Rosenblum EB, Römpler H, Schöneberg T, Hoekstra HE. 2010. Molecular and functional basis of phenotypic convergence in white lizards at White Sands. Proc Natl Acad Sci USA. 107(5):2113–2117. doi:10.1073/pnas.0911042107.

Sheppard PM, Ford EB. 1966. Natural selection and the evolution of dominance. Heredity. 21(1):139–147. doi:10.1038/hdy.1966.8.

Spencer HG, Priest NK. 2016. The evolution of sex-specific dominance in response to sexually antagonistic selection. Am Nat. 187(5):658–666. doi:10.1086/685827.

Stewart AJA, Lees DR. 1987. Genetic control of colour polymorphism in spittlebugs (Philaenus spumarius) differs between isolated populations. Heredity. 59:445–448. doi:10.1038/hdy.1987.154.

Tarutani Y, Shiba H, Iwano M, Kakizaki T, Suzuki G, Watanabe M, Isogai A, Takayama S. 2010. Trans-acting small RNA determines dominance relationships in Brassica self-incompatibility. Nature. 466(7309):983–986. doi:10.1038/nature09308.

West DA. 1977. Melanism in Biston (Lepidoptera: Geometridae) in the rural central Appalachians. Heredity. 39:75–81. doi:10.1038/hdy.1977.43.

Wickham H. 2009. ggplot2: Elegant graphics for data analysis. New York (NY): Springer. doi:10.1007/978-0-387-98141-3.

Wright S. 1929. Fisher’s theory of dominance. Am Nat. 63(685):274–279. doi:10.1086/280260.

Yassin A. 2018. Phylogenetic biogeography and classification of the Drosophila montium species group (Diptera: Drosophilidae). Ann Soc Entomol Fr (N.S.). 54(2):167–175. doi:10.1080/00379271.2018.1447853.

Yassin A, Delaney EK, Reddiex AJ, Seher TD, Bastide H, Appleton NC, Lack JB, David JR, Chenoweth SF, Pool JE, et al. 2016. The pdm3 locus is a hotspot for recurrent evolution of female-limited color dimorphism in Drosophila. Curr Biol. 26(18):2412–2422. doi:10.1016/j.cub.2016.07.016.

Yassin A, Suwalski A, Raveloson Ravaomanarivo LH. 2019. Resolving the synonymy and polyphyly of the ‘Drosophila bakoue species complex’ (Diptera: Drosophilidae: ‘D. montium species group’) with descriptions of two new species from Madagascar. Eur J Taxon. 532:1–26. doi:10.5852/ejt.2019.532.

