## Supplemental Figures for "Interspecific differences in second-site modifiers reshape dominance in a Mendelian trait by altering environmental plasticity"

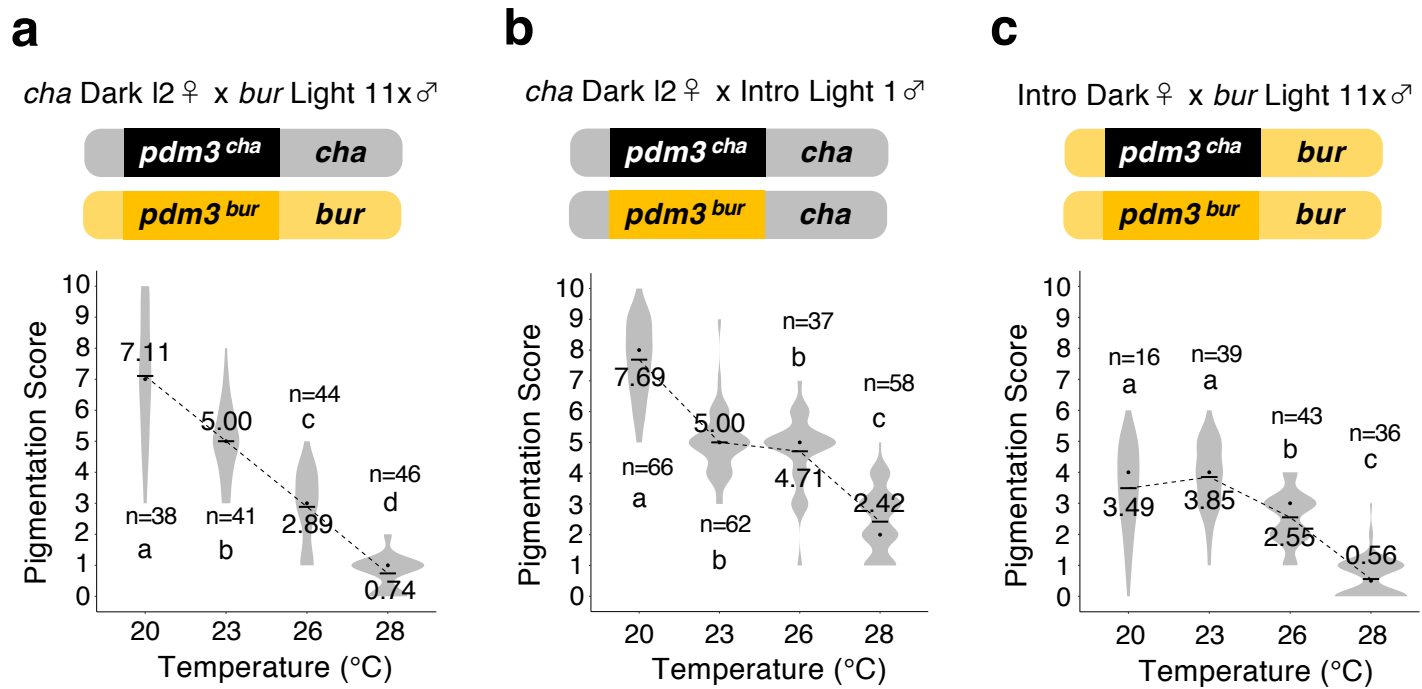

Fig. S1

**a**

*bur* Light 11x ♀ x *bur* Dark 781 ♂

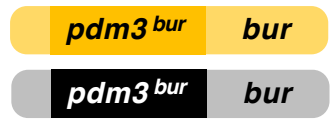

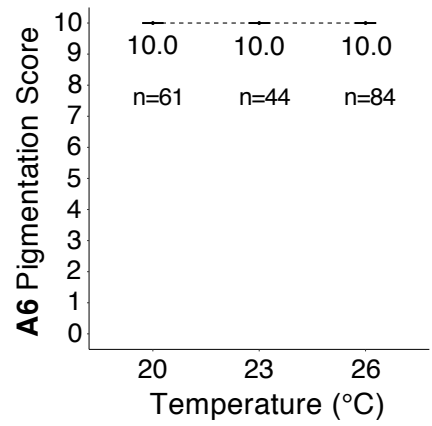

**b**

Intro Light 1 ♀ x *bur* Dark 781 ♂

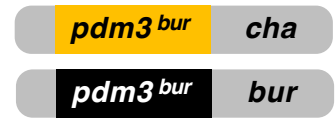

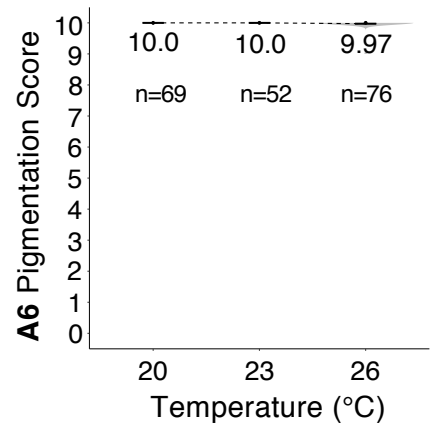

**c**

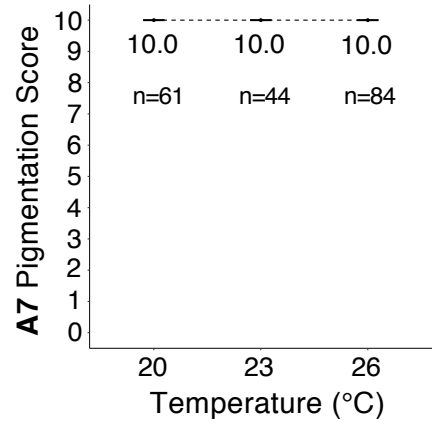

**d**

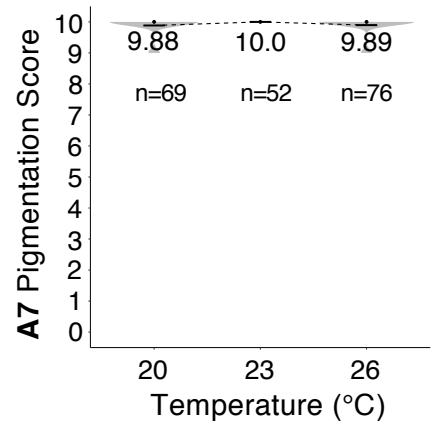

Fig. S2

**a**

*boq* Light 85L ♀ x *boq* Dark I1 ♂

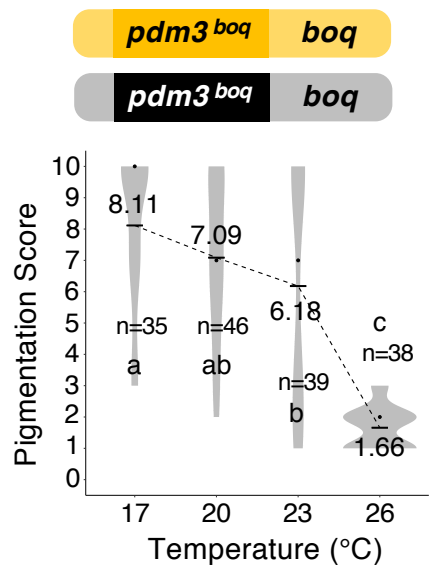

**b**

Intro Light 2 ♀ x *boq* Dark I1 ♂

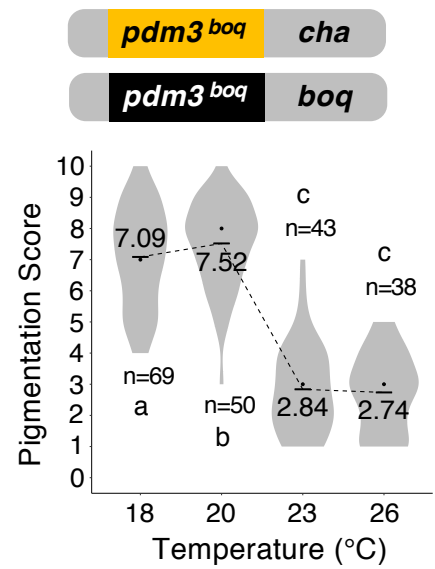

Fig. S3

**a**

*cha* Dark I2 ♀ x *boq* Light 85L ♂

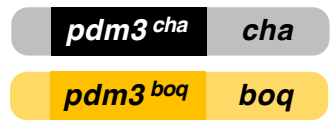

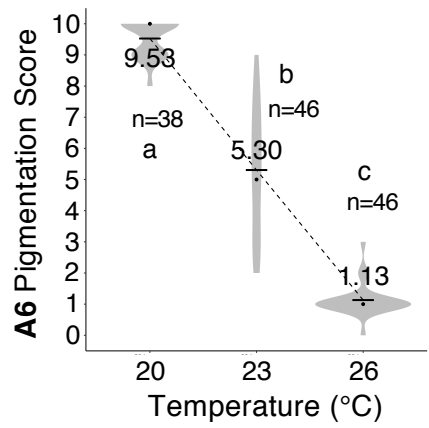

**b**

*cha* Dark I2 ♀ x Intro Light 2 ♂

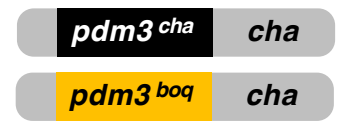

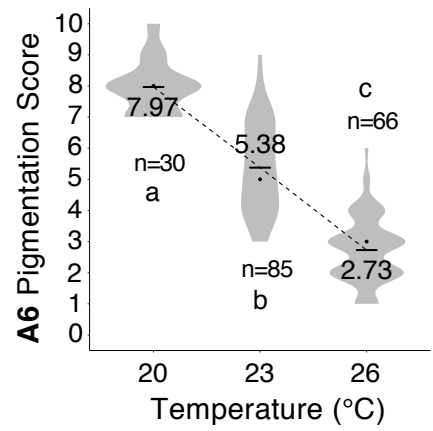

**c**

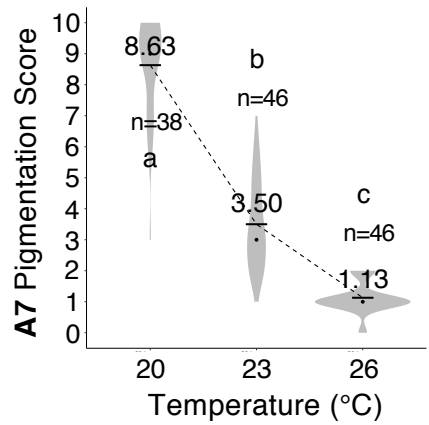

**d**

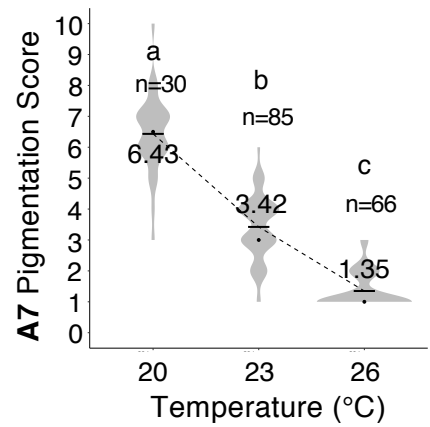

Fig. S4

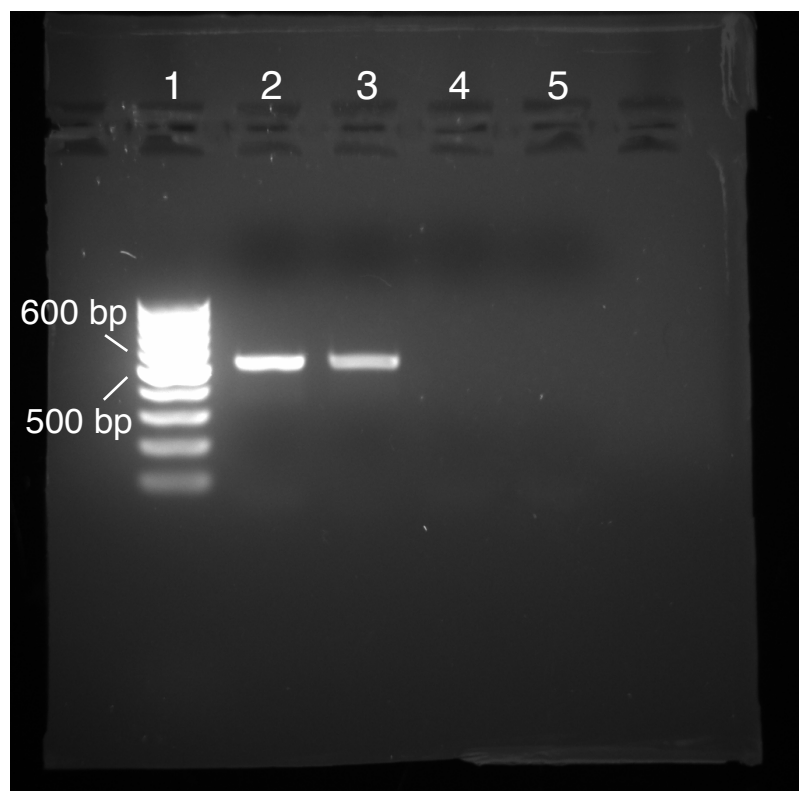

Fig. S5

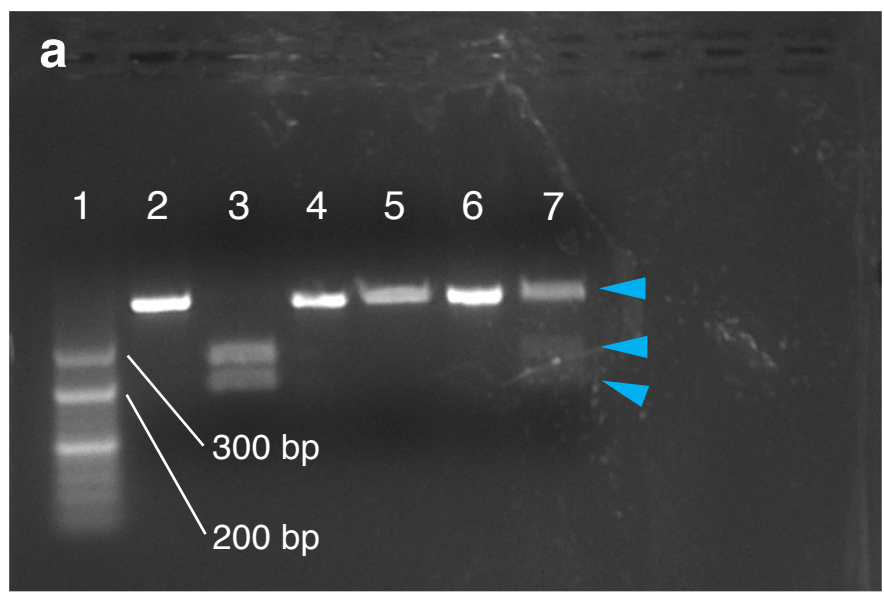

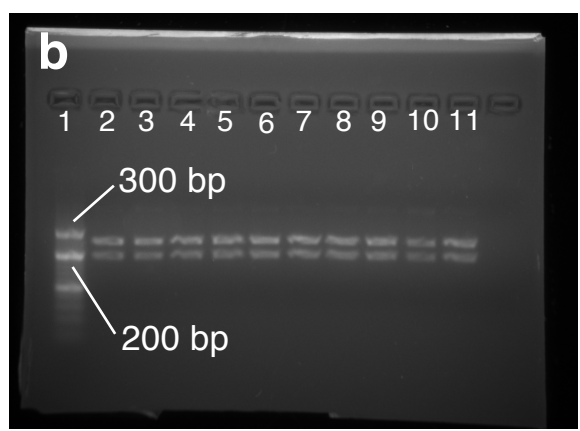

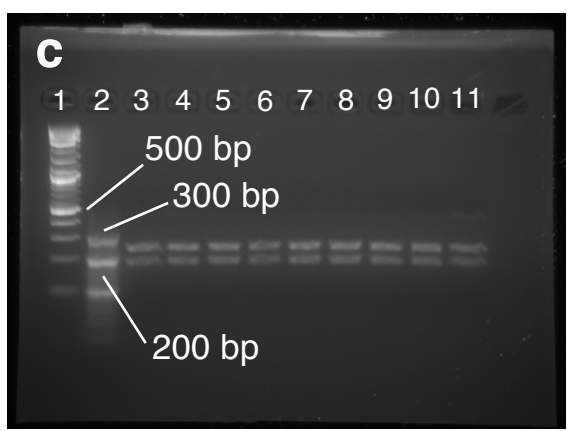

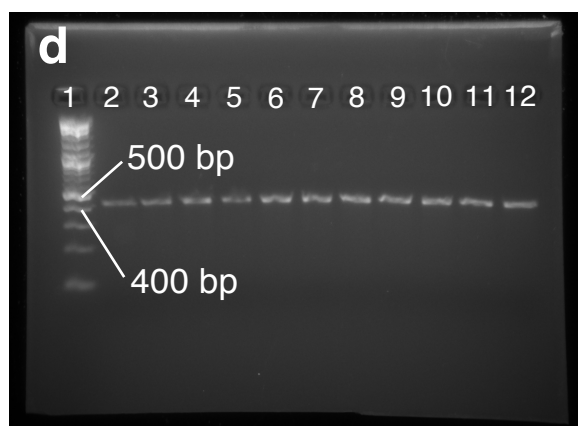

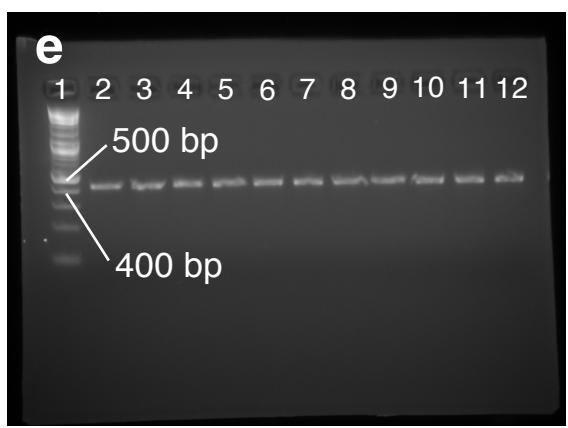

Fig. S6
